# The glutathione import system satisfies the *Staphylococcus aureus* nutrient sulfur requirement and promotes interspecies competition

**DOI:** 10.1101/2021.10.26.465763

**Authors:** Joshua M. Lensmire, Michael R. Wischer, Lo M. Sosinski, Elliot Ensink, Jack P. Dodson, John C. Shook, Phillip C. Delekta, Christopher C. Cooper, Daniel Havlichek, Martha H. Mulks, Sophia Y. Lunt, Janani Ravi, Neal D. Hammer

**Affiliations:** Department of Microbiology and Molecular Genetics, Michigan State University, East Lansing, MI 48824, USA; Department of Pathobiology and Diagnostic Investigation, Michigan State University, East Lansing, MI 48824, USA; Department of Biochemistry and Molecular Biology Michigan State University, East Lansing, MI 48824, USA; Department of Medicine, Division of Infectious Disease, Michigan State University, East Lansing, MI 48824, USA; Department of Chemical Engineering and Materials Science, Michigan State University, East Lansing, MI 48824, USA

## Abstract

Sulfur is an indispensable element for proliferation of bacterial pathogens. Prior studies indicated that the human pathogen, *Staphylococcus aureus* utilizes glutathione (GSH) as a source of nutrient sulfur; however, mechanisms of GSH acquisition are not defined. Here, we identify a previously uncharacterized five-gene locus comprising a putative ABC-transporter and γ–glutamyl transpeptidase (*ggt*) that promotes *S. aureus* proliferation in medium supplemented with either reduced or oxidized GSH (GSSG) as the sole source of nutrient sulfur. Based on these phenotypes, we name this transporter the **<u>G</u>**lutathione **<u>i</u>**mport **<u>s</u>**ystem (GisABCD). We confirm that Ggt is capable of cleaving GSH and GSSG γ–bonds and that this process is required for their use as nutrient sulfur sources. Additionally, we find that the enzyme is cell associated. Bioinformatic analyses reveal that only *Staphylococcus* species closely related to *S. aureus* encode GisABCD-Ggt homologues. Homologues are not detected in *Staphylococcus epidermidis*. Consequently, we establish that GisABCD-Ggt provides a competitive advantage for *S. aureus* over *S. epidermidis* in a GSH-dependent manner. Overall, this study describes the discovery of a nutrient sulfur acquisition system in *S. aureus* that targets GSH and promotes competition against other staphylococci commonly associated with the human microbiota.

---

Bacterial pathogens are limited to metabolites present in host tissues to fulfill nutritional requirements during infection. Strategies pathogens employ to acquire nutrient transition metals, such as iron, are well established^1,2^. However, studies defining mechanisms that support nutrient sulfur acquisition have been restricted to a limited number of pathogens^3^. Sulfur is essential due to its capacity to fluctuate between redox states and therefore catalyzes numerous cellular reactions^4,5^. Ultimately, cells require sulfur to synthesize cysteine (Cys). Cys is the fulcrum of sulfur metabolism as an intermediate for methionine (Met) and cofactors such as Fe-S clusters^4,6–8^. In host cells and some bacterial species, Cys is also required to generate the low molecular weight thiol glutathione (GSH)^9^.

GSH concentrations range between 0.5 and 10 mM in mammalian tissues, making it a relatively abundant source of nutrient sulfur for invading pathogens^9–20^. In addition to Cys, GSH consists of glutamate and glycine. A unique γ-bond links the glutamate γ-carboxyl to the Cys amine. To maintain Cys reservoirs, organisms rely on GSH catabolism via the γ-glutamyl cycle^9,21^. Liberation of Cys from GSH is a two-step process that requires γ-glutamyl transpeptidase (Ggt), a specialized peptidase conserved in all domains of life due to its role in the γ-glutamyl cycle^9,22–25^. Typically, Ggt is localized to the periphery of the cell where it degrades endogenous GSH in the extracellular milieu or within the Gram-negative periplasm^26–28^. Ggt has been implicated in pathogen nutrient sulfur acquisition via Cys liberation in only one species, *Francisella tularensis*^18^.

*Staphylococcus aureus* is the leading cause of superficial and invasive bacterial diseases in the United States and Europe^29–31^. Strategies *S. aureus* employs to obtain nutrient sulfur during pathogenesis are largely unknown. Moreover, while GSH is an established *in vitro* source of nutrient sulfur for *S. aureus*, mechanisms of GSH acquisition have not been defined^32^. *S. aureus* does not synthesize GSH but encodes a putative Ggt. This fact supports the hypothesis that staphylococcal Ggt catabolizes exogenous host GSH, liberating Cys as a means to satisfy the nutrient sulfur requirement ^33^. Here we demonstrate that *S. aureus* utilizes oxidized GSH (GSSG) as a nutrient sulfur source and isolate mutants that fail to proliferate in medium supplemented with GSSG or GSH as the sole source of nutrient sulfur. These mutants harbored transposon (Tn) insertions within a five gene locus, SAUSA300_0200-0204. SAUSA300_0200-0203 encodes a predicted ATP-binding-cassette (ABC) transporter and SAUSA300_0204 encodes a putative Ggt. Consequently, we name this transporter the **<u>G</u>**lutathione **<u>i</u>**mport **<u>s</u>**ystem (GisABCD). We determine that *S. aureus* Ggt associates with the cell and that the recombinant enzyme cleaves both GSH and GSSG. A query for GisABCD-Ggt across Firmicutes revealed that only a select clade within the *Staphylococcus* genus, one that does not include *S. epidermidis*, encodes homologues of the system. Consistent with this, *S. aureus* outcompetes *S. epidermidis* in GSSG- or GSH-supplemented conditions. Therefore, this newly described nutrient sulfur acquisition system provides a competitive advantage for *S. aureus* over other staphylococci commonly associated with the human microbiota.

## Results

### *S. aureus* proliferates in medium supplemented with GSSG as the sole source of nutrient sulfur

A previous study qualitatively reported that *S. aureus* generates colonies on a chemically-defined medium supplemented with GSH as the sole sulfur source, indicating that the abundant host metabolite is a viable source of nutrient sulfur^32^. However, *S. aureus* likely encounters both reduced and oxidized GSH (GSSG) as the pathogen induces a potent oxidative burst during infection^34^. Therefore, we hypothesized that *S. aureus* utilizes GSSG as a source of nutrient sulfur. To quantitatively assess utilization of GSH and GSSG as sulfur sources by *S. aureus*, a chemically-defined medium, referred to as PN, was employed^35^. PN contains sulfate and methionine (Met), but *S. aureus* lacks the capacity to assimilate sulfate or utilize Met as sources of sulfur; thus, cystine (oxidized cysteine or CSSC) is added to fulfill the sulfur requirement^32,36^. In keeping with this, a USA300 LAC strain of *S. aureus* (JE2) fails to proliferate in PN devoid of CSSC (**Fig. 1a**). Notably, replacing CSSC with either 50 μM GSH or 25 μM GSSG stimulates *S. aureus* proliferation (**Fig. 1a**). *S. aureus* utilization of GSSG expands the number of sulfur-containing metabolites present in host tissues that support proliferation of this pathogen. To determine whether utilization of GSSG is conserved throughout the species, we examined proliferation of clinical isolates in GSSG-supplemented medium. Growth of methicillin-sensitive (MSSA) and methicillin-resistant (MRSA) clinical isolates was quantified in PN supplemented with GSSG as the sole sulfur source. Compared to PN lacking a viable sulfur source, GSSG supplementation substantially increased terminal OD_600_ (**Fig. 1b and Supplementary Fig. S1**). Additionally, GSSG supplementation also considerably increased the terminal OD_600_ of other *S. aureus* strains (**Fig. 1c and Supplementary Fig. S1**).

**Figure 1.**
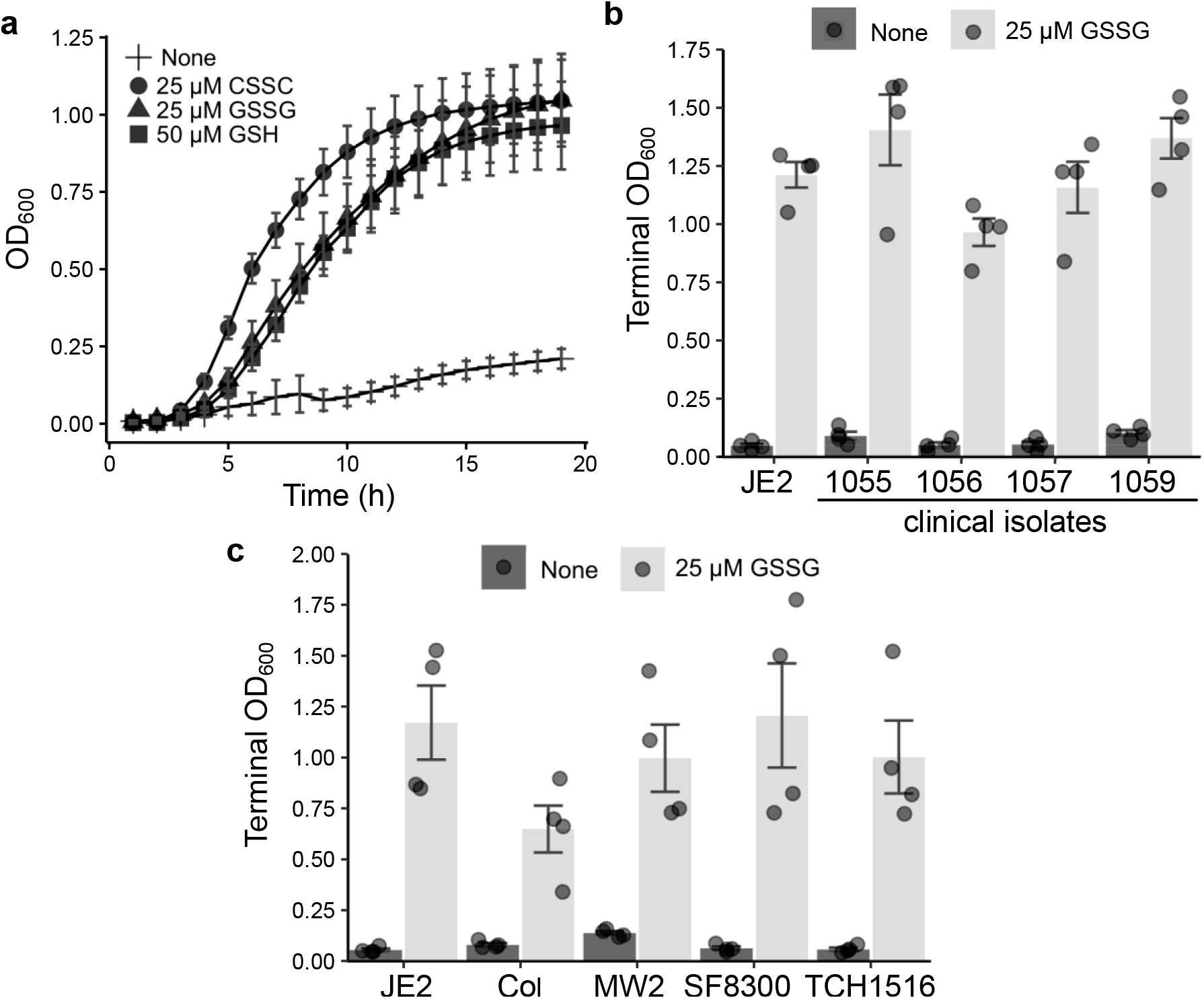
Supplementation of GSSG as the sole source of nutrient sulfur supports proliferation of *S. aureus*. **a**, WT JE2 *S. aureus* was cultured in PN medium lacking a viable sulfur source. CSSC, GSSG, or GSH were added to the medium at indicated concentrations. **b**, JE2 and either MSSA or MRSA clinical isolates were grown in PN supplemented without a viable sulfur source (None) or 25 μM GSSG for 19 hours (h). **c**, *S. aureus* strains were cultured in PN medium supplemented without sulfur (None) or 25 μM GSSG and cultured for 25 h. Bars depict the mean terminal optical density at 600 nm (OD_600_), and circles represent individual replicate terminal OD_600_. The mean of at least three independent trials and error bars representing ± 1 standard error of the mean are presented in **a, b**, and **c**.

### The SAUSA300_0200-0204 locus supports *S. aureus* utilization of GSH and GSSG as sulfur sources

To determine genetic factors required for *S. aureus* utilization of GSSG as a sulfur source, we screened the Nebraska Transposon Mutant Library for mutants that fail to proliferate in PN medium supplemented with 25 μM GSSG as the sole source of sulfur^37^. Five GSSG proliferation-impaired mutants were identified in the screen, each harboring an independent transposon insertion in one of five genes present in the SAUSA300_0200-*ggt* locus (**Fig. 2a**). Notably, SAUSA300_0200-0203 encodes a putative nickel-peptide ABC transporter. Backcrossing transposon-inactivated genes into an otherwise wild type (WT), isogenic JE2 strain significantly decreased proliferation in medium supplemented with 25 μM GSSG (**Fig. 2b**). The transposon mutants also displayed decreased proliferation in PN supplemented with 50 μM GSH as the sole sulfur source (**Fig. 2b**). However, the backcrossed mutant strains demonstrated WT-like growth in medium supplemented with 25 μM CSSC or in a rich medium, indicating the proliferation defect is specific to GSH and GSSG (**Fig. 2b**). To address complications associated with auto-oxidation of GSH to GSSG in aerobic environments, we tested anaerobic proliferation in media supplemented with GSH or GSSG. Mutant strains harboring a transposon in SAUSA300_0201 or an in-frame deletion of all five genes were used in this assessment. In these conditions, both mutant strains proliferate to WT levels in PN supplemented with CSSC but display reduced growth upon GSH or GSSG supplementation (**Supplementary Fig. S2**). This finding confirms that SAUSA300_0200-*ggt* supports *S. aureus* utilization of both GSH and GSSG as distinct sources of nutrient sulfur. Complementation experiments tested whether proliferation of a *ggt* mutant cultured in PN medium supplemented with GSH or GSSG could be restored by providing WT or a C-terminal His-tagged *ggt* encoded on a plasmid. *ggt* mutant strains harboring either plasmid exhibit WT-like growth, confirming that failure to proliferate in GSSG- or GSH-supplemented medium is due to genetic inactivation of *ggt* (**Supplementary Fig. S3**).

**Figure 2.**
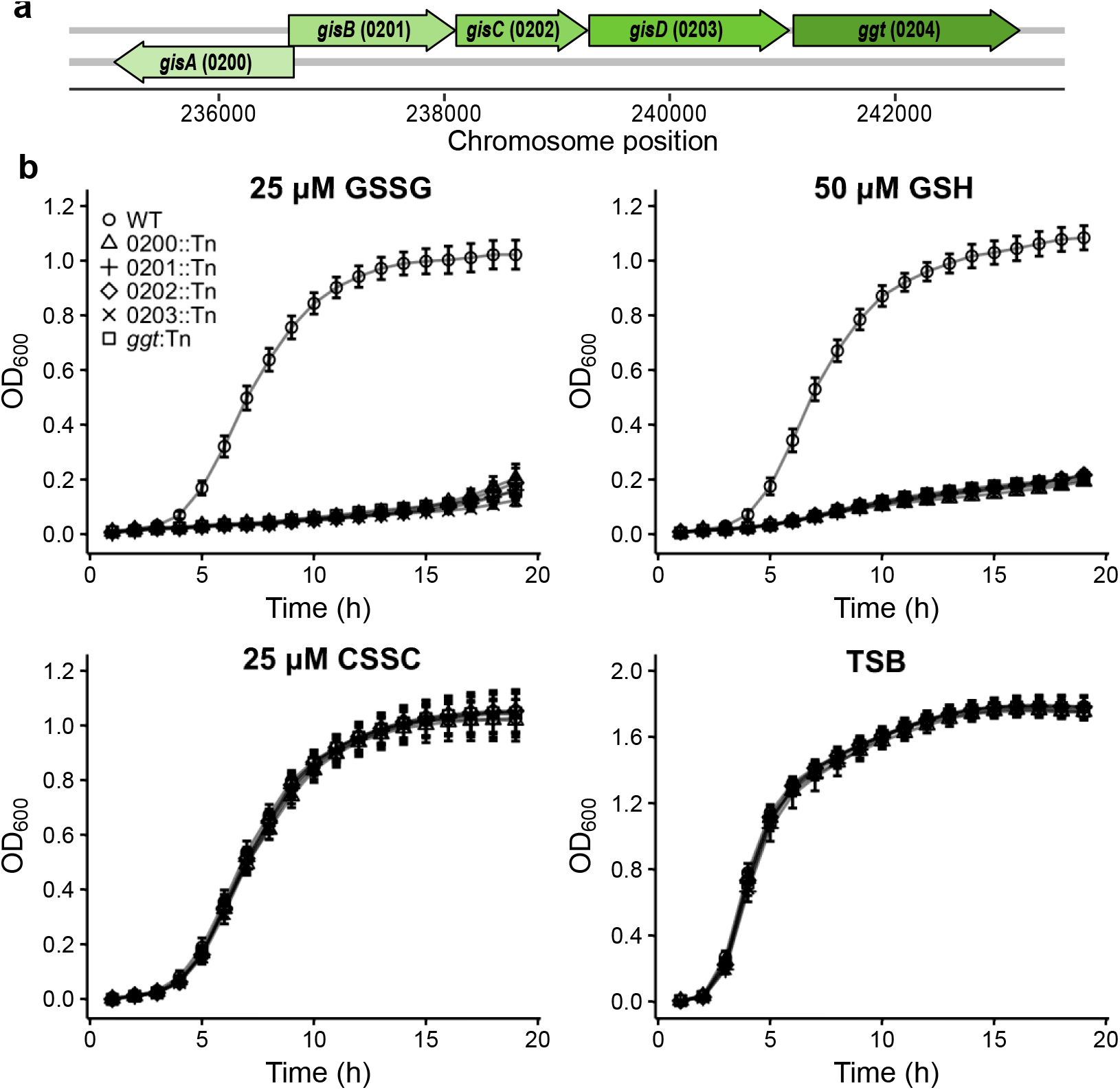
SAUSA300_0200-*ggt* supports *S. aureus* proliferation on GSH and GSSG as sources of nutrient sulfur. **a**, Orientation of SAUSA300_0200-*ggt* within the *S. aureus* genome. **b**, Strains cultured in tryptic soy broth (TSB) or PN supplemented with 25 μM GSSG, 50 μM GSH, or 25 μM CSSC. The mean of at least three independent trials and error bars representing ± 1 standard error of the mean are presented.

Based on the inability of the mutant strains to proliferate in GSH- or GSSG-supplemented PN medium and the fact that SAUSA300_0200-SAUSA300_0203 encodes a putative ABC-transporter with a predicted Ggt (SAUSA300_0204), we named these genes the **<u>g</u>**lutathione **<u>i</u>**mport **<u>s</u>**ystem (*gisABCD-ggt*) (**Fig. 2a**). Domain architecture analysis of the GisABCD-Ggt system reveals that GisA contains ATP-binding cassette domains (**Supplementary Fig. S4**). In support of the annotation, recombinant GisA purified from *Escherichia coli* demonstrates ATP hydrolysis activity (**Supplementary Fig. S5**). GisB and GisC are transmembrane permeases with nine predicted transmembrane segments while GisD contains a signal peptide with a putative lipid attachment site (**Supplementary Fig. S4**). Pfam analysis predicts that Ggt is a γ-glutamyl transpeptidase (**Supplementary Fig. S4**).

### Ggt hydrolyzes GSH and GSSG γ-bonds and is cell associated

A hallmark of γ-glutamyl transpeptidase is its capacity to cleave the GSH γ–bond, liberating glutamate. To validate the Pfam domain prediction, we sought to determine whether *S. aureus* Ggt cleaves the γ–bond linking glutamate to Cys in GSH and GSSG ^38,39^. C-terminal His-tagged recombinant Ggt (rGgt) was expressed and purified from *E. coli* (**Supplementary Fig. S6**). In other species, Ggt enzymes are translated as an inactive polypeptide that is auto-catalytically cleaved to generate approximate 40 kDa and 35 kDa subunits^39,40^. Consistent with this, a tripartite banding pattern consisting of full-length pro-Ggt (75 kDa) and smaller, mature enzyme subunits (40 kDa and 35 kDa) are observed (**Supplementary Fig. S6**). Mature Ggt cleaves GSH by attacking the glutamyl residue, transferring it to the enzyme. Ultimately, water hydrolyzes the γ-bond, liberating glutamate^41^. Therefore, to quantify *S. aureus* Ggt γ-glutamyl transpeptidase activity, rGgt was incubated with increasing concentrations of GSH or GSSG and glutamate release was measured by mass spectrometry. Glutamate was detected in reactions containing rGgt incubated in the presence of either GSH or GSSG (**Fig. 3a**). Importantly, glutamate was not detected in reactions lacking substrate or rGgt, indicating glutamate release resulted from enzymatic activity (data not shown). K_m_ values of Ggt for GSSG and GSH were determined to be 39 μM and 58.5 μM, respectively. These values are similar to previously reported Ggt homologues expressed in other organisms. For example, the *E. coli* Ggt GSH K_m_ value is 29 μM^42^. These data support the *in silico* prediction and provide a molecular explanation for the *ggt* mutant proliferation defect in medium supplemented with GSH or GSSG as sources of nutrient sulfur—*ggt* mutant cells are unable to initiate liberation of Cys due to an inability to hydrolyze the GSH or GSSG γ-bond. Next, we attempted to determine whether hydrolysis of GSH and GSSG occurs intracellularly or extracellularly by localizing Ggt to *S. aureus* subcellular fractions.

**Figure 3.**
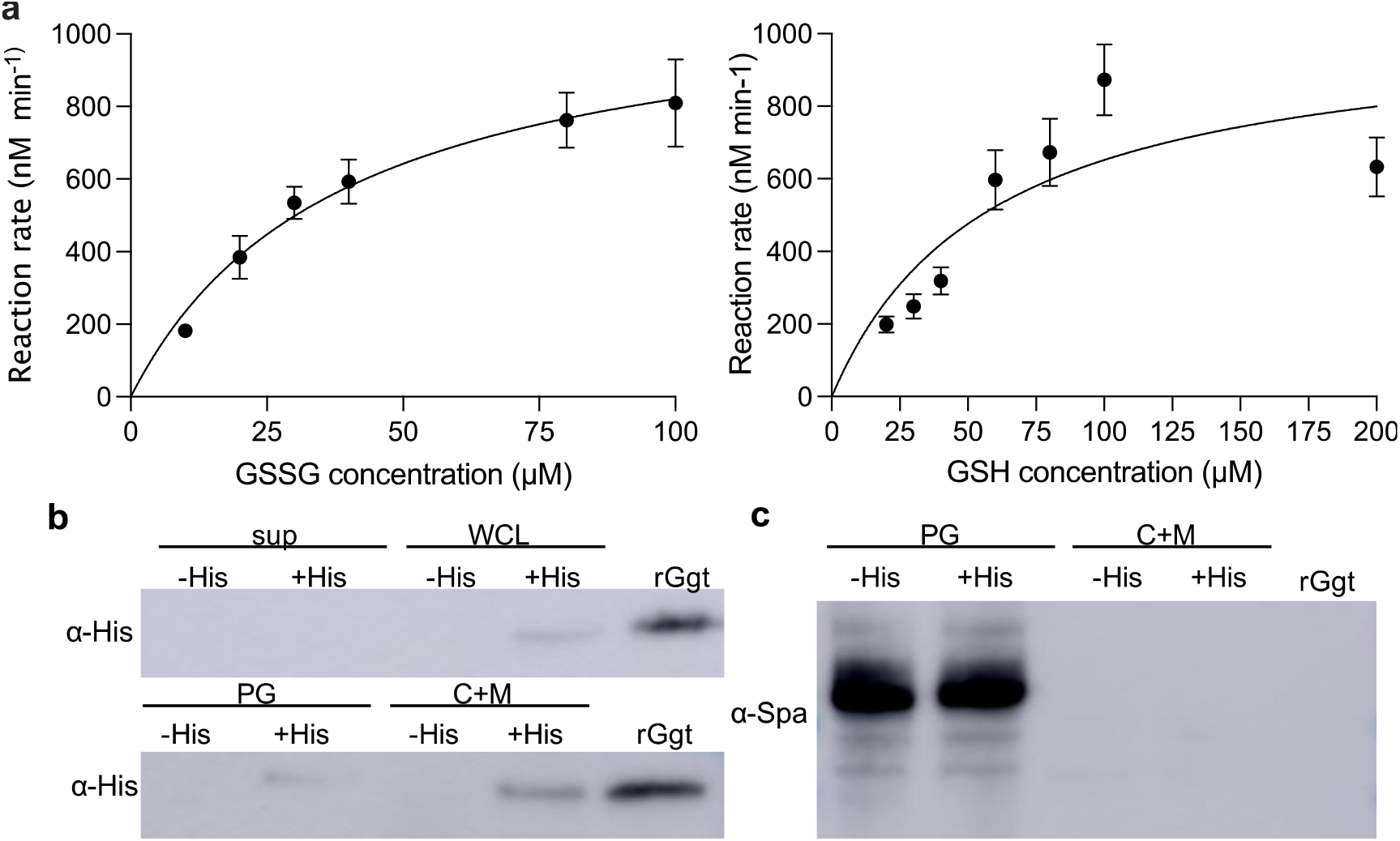
*S. aureus* Ggt liberates glutamate from GSH and GSSG and is cell-associated. **a**, rGgt was incubated in the presence of indicated GSH and GSSG concentrations. Mean glutamate release per minute was measured using four independent rGgt protein preparations. Glutamate released per min was calculated and the data was fit with the Michaelis Menten equation using GraphPad Prism. Error bars represent ± standard error of the mean. **b and c**, Fractions of *S. aureus ggt*∷Tn harboring pOS1 P_*lgt*_∷*ggt* (-His) and pOS1 P_*lgt*_∷*ggt*-His (+His) probed with α-His antibodies. Fractions include the culture supernatant (sup), whole cell lysate (WCL), peptidoglycan (PG), and protoplast lysate which contains the cytoplasm and membranes (C+M). **c**, Fractions were probed with anti-protein A (Spa) antibodies (α-Spa).

*Bacillus sp*. secrete Ggt; however, structural predictions of *S. aureus* Ggt failed to detect canonical secretion signal sequences within the primary sequence (**Supplementary Fig. S4**). For example, SignalP-5.0 predictions detected considerably low likelihoods of signal peptide, twin-arginine translocation (TAT) signal peptide, or lipoprotein signal peptide sequences (0.007, 0.003, 0.008, respectively). TatP 1.0 also failed to predict a signal peptide^43,44^. Ggt localization varies depending on the organism, but here it has implications for the substrate of GisABCD. Extracellular Ggt supports a model whereby GisABCD imports Ggt cleavage products, whereas intracellular Ggt suggests GisABCD imports GSH and GSSG intact^18,19^. We used the previously described His-tagged Ggt expression vector (Ggt-His) that functionally complements the *ggt*∷Tn mutant (**Supplementary Fig. S3**) to determine subcellular localization of the enzyme. *S. aureus* cells expressing native or Ggt-His were cultured in PN supplemented with 25 μM GSSG, collected, and fractionated into supernatant and whole cell lysate (WCL) samples. An α-6xHis antibody was used to monitor Ggt-His within each fraction. rGgt served as a size comparison control. Ggt-His was not detected in the supernatant fractions; however, a band at ~35kDa was observed in both the Ggt-His WCL and rGgt samples. This band is specific to Ggt-His as it was not observed in WCL generated from cells expressing Ggt lacking the His-tag and corresponds to the rGgt subunit containing the His-tag. Presence of Ggt-His signal within the WCL fraction supports the conclusion that Ggt is cell-associated (**Fig. 3b**). WCL was further fractioned by removal of the peptidoglycan (PG). Resulting protoplasts were lysed to generate a fraction containing cytoplasm and membranes (C+M). His-dependent ~35 kDa signal was increasingly apparent in the C+M fraction compared to the PG fraction when equivalent levels of total protein are assessed (**Fig. 3b and 3c**). Overall, the lack of a secretion signal sequence and enrichment in the C+M fraction support a model whereby GSH and GSSG are imported intact and catabolized in cytoplasm, leading to the eventual liberation of Cys to satisfy the nutrient sulfur requirement (**Supplementary Fig. S7**). Cytoplasmic localization of Ggt has been reported in only one other bacterial pathogen, *Neisseria meningitidis*^28^. Given that *S. aureus* does not synthesize GSH and therefore does not utilize GSH as its major low molecular redox thiol, we predict that intracellular Ggt will be associated with other bacterial species that lack GSH synthesis. Additionally, we surmise that it will function primarily to satisfy the demand for nutrient sulfur in these species^45,46^.

### GisABCD-Ggt is not required for systemic infection of *S. aureus*

We next tested whether GisABCD-Ggt is important for *S. aureus* pathogenesis using a systemic murine model of infection. However, organs of mice infected with a *gisB∷*Tn mutant contained equivalent bacterial burdens compared to WT-infected animals (**Supplementary Fig. S8**). This result indicates that GisABCD-Ggt is dispensable for *S. aureus* pathogenesis during systemic infection. A possible explanation for the lack of a virulence defect is that *S. aureus* may encode multiple GSH and GSSG transporters. In fact, while GisABCD-Ggt supports proliferation of *S. aureus* in micromolar concentrations of GSSG or GSH, increasing GSH concentrations closer to those present in host tissues restores Δ*gisABCD-ggt* mutant proliferation in PN medium (**Supplementary Fig. S9**). Increasing GSSG concentrations did not stimulate Δ*gisABCD-ggt* mutant proliferation. These results support the conclusion that GisABCD-Ggt is an absolute requirement for GSSG utilization but that another, potentially low affinity, GSH transporter is also active in this pathogen.

### *S. aureus* GisABCD-Ggt is conserved in select Firmicutes

In an attempt to identify a function for GisABCD-Ggt beyond systemic host colonization, we quantified conservation of the system throughout Firmicutes. Due to the ubiquity of ABC-transporters across bacterial genera, we first focused on potential Ggt homologues. We found that numerous Firmicutes encode Ggt homologues, including *Bacillus, Gracilibacillus, Lysinibacillus*, and *Brevibacterium*; however, subsequent identification of potential GisABCD homologues was limited to a minority of these bacteria (**Supplementary Fig. S10**). Overall distribution of GisABCD-Ggt homologues across Firmicutes revealed six distinct clusters. Cluster 3 is the least populated, as species in this cluster encode only Ggt homologues. Firmicutes in clusters 2 and 4 contain Ggt and either a GisB or a GisC homologue, respectively. Cluster 5 contains bacteria that encode GisB, GisC, and Ggt homologues (**Supplementary Fig. S10**). Cluster 6 includes a few bacilli species that encode proteins similar to the complete *S. aureus* GisABCD-Ggt system, but Cluster 1 stands apart as it encompasses the *S. aureus* query sequences and comprises species that harbor complete GisABCD-Ggt systems at greater than >80% similarity. Notably, this cluster is distinct from other Firmicutes and is restricted to members of the *Staphylococcus* genus with putative operons encoding proteins most similar to *S. aureus* GisABCD-Ggt (**Supplementary Fig. S10**).

We predicted that GisABCD-Ggt would be widespread throughout the genus *Staphylococcus*. Yet, only species in the *S. aureus*-related cluster complex—which includes *Staphylococcus argenteus, Staphylococcus schweitzeri*, and *Staphylococcus simiae—*encode homologues of a complete GisABCD-Ggt system (**Fig. 4a**)^47,48^. Conservation rapidly diverged as the next related species, *Staphylococcus epidermidis*, encodes apparent GisA, GisC, and Ggt homologues with exceedingly low percent similarities (**Fig. 4a**)^48^. Lack of GisABCD-Ggt conservation supports the hypothesis that *S. epidermidis* is incapable of utilizing low concentrations of GSH as a source of nutrient sulfur.

**Figure 4.**
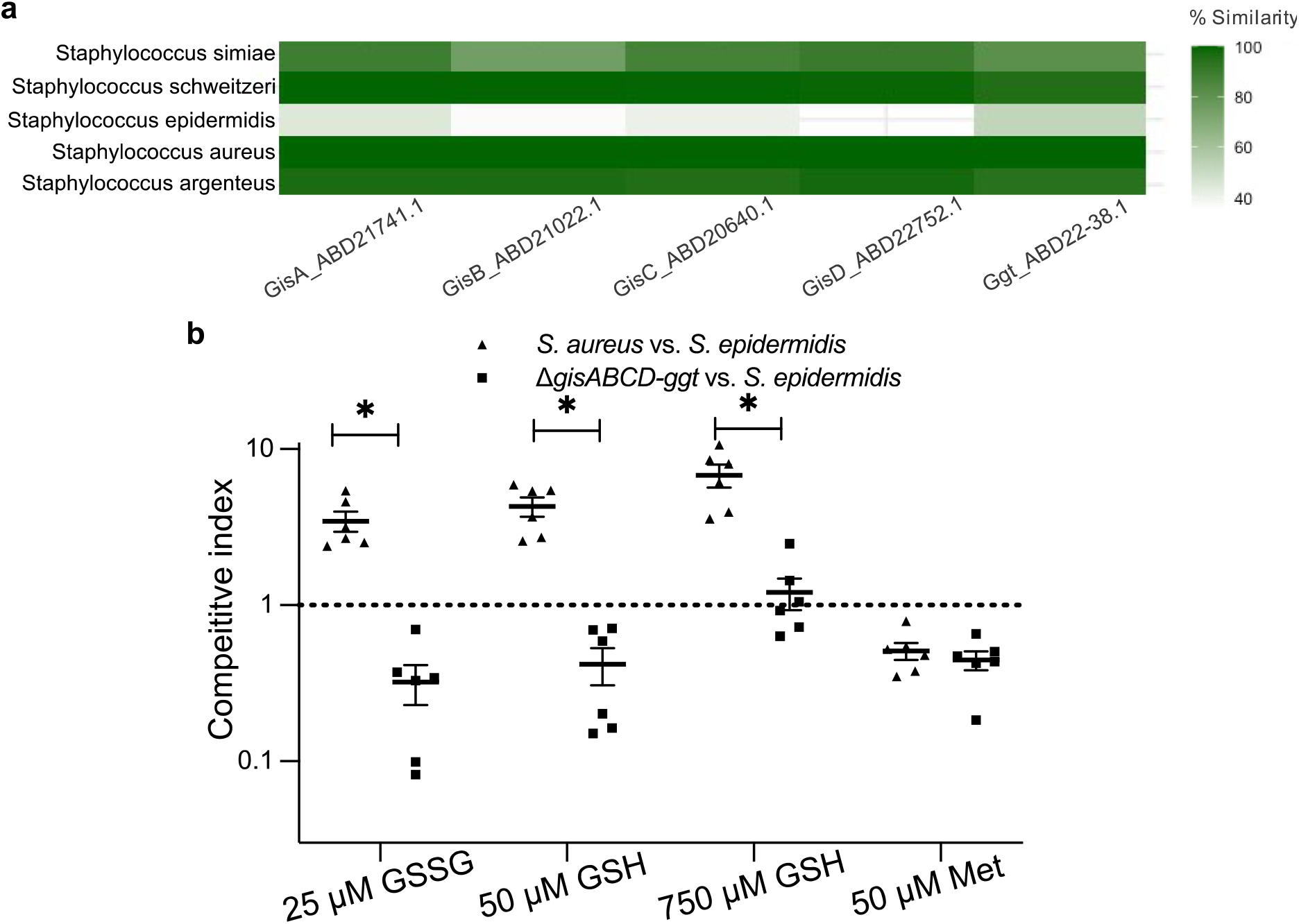
GisABCD is conserved in exclusive staphylococci and promotes competition in GSH- or GSSG-supplemented media. **a**, Heatmap depicting percent similarities of GisABCD-Ggt proteins across Staphylococci (*S. aureus* Gis-Ggt proteins were used as the starting points for the homology searches). **b**, *In vitro* competition experiments between *S. aureus* and *S. epidermidis* strain RP62a. The line represents the mean competitive index for each individual trial. Error bars represent ± 1 standard error of the mean. * Indicates p-value <0.05 as determined by one-way ANOVA with Tukey’s multiple test correction.

### GisABCD-Ggt promotes interspecies *Staphylococcus* competition in a GSH-specific manner

To quantify *S. epidermidis* sulfur source dependent proliferation, we first needed to define whether the organism is capable of utilizing Met, a component of PN medium, as a source of nutrient sulfur. Additionally, *S. epidermidis* encodes predicted sulfate assimilation enzymes; thus, sulfate (MgSO_4_) was removed from PN medium^49,50^. Supplementation of sulfate-depleted PN with Met as the sole source of nutrient sulfur stimulates proliferation of *S. epidermidis*, but not *S. aureus* (**Supplementary Fig. S11a**). Therefore, Met was also removed and the resulting medium, PN_mod_, was used to quantify GSH- and GSSG-dependent proliferation of *S. epidermidis* compared to *S. aureus*. Consistent with the importance of GisABCD-Ggt to *S. aureus* GSSG utilization, supplementation of 25 μM GSSG failed to stimulate appreciable proliferation of *S. epidermidis*. Supplementation with 50 μM GSH promoted delayed growth of *S. epidermidis* relative to *S. aureus* (**Supplementary Fig. S11a**). Increasing GSH concentrations to 750 μM resulted in comparable proliferation between *S. epidermidis* and *S. aureus* (**Supplementary Fig. S11a**). These data indicate potential conservation of the unknown GSH acquisition system between the two species. *S. epidermidis* does not exhibit enhanced growth in 375 μM GSSG relative to 25 μM GSSG (**Supplementary Fig. S11a**), a result similar to the *S. aureus* Δ*gisABCD-ggt* mutant and consistent with the conclusion that GisABCD-Ggt is an absolute requirement for GSSG utilization in *Staphylococcus*.

Next, the capacity of GisABCD-Ggt to provide a competitive advantage to *S. aureus* over *S. epidermidis* was determined. To test this, output CFU ratios of *S. aureus* to *S. epidermidis* after a 24 h co-culture in PN_mod_ supplemented with different sulfur sources were quantified. *S. aureus* outcompeted both a *S. epidermidis* clinical isolate and the laboratory RP62a strain in PN_mod_ supplemented with 25 μM GSSG or 50 μM GSH (**Fig. 4b**, **Supplementary Fig. S11b**). Conversely, *S. epidermidis* strains outcompeted *S. aureus* in medium with 50 μM Met (**Fig. 4b**, **Supplementary Fig. S11b**). *S. aureus* exhibited a competitive advantage over *S. epidermidis* in 750 μM GSH, despite equivalent *S. epidermidis* proliferation in monoculture at this concentration (**Fig. 4b, Supplementary Fig. S11b**). Both *S. epidermidis* strains outcompeted *S. aureus ΔgisABCD-ggt* in 25 μM GSSG and 50 μM GSH, underscoring the importance of GisABCD-Ggt in promoting *S. aureus* competition over *S. epidermidis* in environments containing limiting concentrations of GSH or GSSG. Equivalent quantities of *S. epidermidis* and *S. aureus ΔgisABCD-ggt* were recovered in medium supplemented with 750 μM GSH (**Fig. 4b, Supplementary Fig. S11b**). These findings provide additional support to the conclusion that Gis-independent GSH acquisition is conserved between *S. aureus* and *S. epidermidis*.

## Discussion

This study expands the number of host-derived metabolites that satisfy the *S. aureus* nutrient sulfur requirement to include GSSG. Our results demonstrate that the ABC-transporter system, GisABCD, and γ-glutamyl transpeptidase, Ggt, support acquisition of GSH and GSSG as sources of nutrient sulfur. We provide evidence that Ggt is cell associated. Cytoplasmic localization of Ggt would be notable given that *S. aureus* does not utilize GSH as a low molecular weight thiol and therefore suggests that the sole function of Ggt is nutrient sulfur catabolism. However, our localization studies do not rule out the possibility that Ggt is anchored to the outer leaflet of the plasma membrane. This localization pattern would support a model whereby GSH or GSSG cleavage occurs first and is followed by transport of the Cys-Gly product through GisABCD. Additional studies are required to corroborate the lack of canonical secretion sequences and confirm intracellular Ggt localization. Nonetheless, we show that GisABCD-Ggt is tuned to capture and catabolize exogenous GSH and GSSG for use as sources of nutrient sulfur.

Failure of the *gisB*∷Tn mutant to exhibit an overt virulence defect is partially explained by the finding that *in vitro* supplementation with GSH concentrations mimicking those present in host tissues stimulates *gis* mutant proliferation. This result is consistent with the fact that the host invests considerable energy preserving redox homeostasis by maintaining GSH quantities in vast excess to GSSG^9,51^. Therefore, we demonstrate that while *S. aureus* utilizes the specialized GisABCD-Ggt system to scavenge GSH and limiting GSSG, the pathogen has also evolved multiple strategies to target the more abundant GSH.

Conservation of GisABCD-Ggt revealed the system is encoded in an exclusive clade of *Staphylococcus* species that includes *S. argenteus, S. schweitzeri*, and *S. simiae*, but not *S. epidermidis*. The fact that *S. epidermidis* proliferates poorly in a GSSG-supplemented medium and that GisABCD-Ggt promotes *S. aureus* competition over *S. epidermidis* in GSH- and GSSG-limiting environments further underscores the importance of GisABCD-Ggt to acquisition of GSSG and GSH as sources of nutrient sulfur. *S. argenteus* colonizes fruit bats and monkeys and causes skin and soft tissue infections and sepsis in humans^52–59^. *S. epidermidis*, *S. aureus*, and *S. schweitzeri* colonize nasal passages of humans or closely related primates, but the lack of GisABCD-Ggt conservation in *S. epidermidis* implies the system is not essential for nasal colonization^47,60–62^. However, both *S. aureus* and *S. epidermidis* are also common residents of the human skin microflora. Therefore, this environment could be further explored to determine whether GisABCD-Ggt provides a competitive advantage to *S. aureus* and *S. argenteus* over *S. epidermidis* in this dynamic host niche.

## Materials and Methods

### Bacterial strains used in this study

The WT *S. aureus* strain used in these studies was JE2, a laboratory derivative from the community-acquired, methicillin-resistant USA300 LAC^63^. SAUSA300_0200*-ggt* mutant strains were generated via transduction of the transposon-inactivated gene from the Nebraska Transposon Mutant Library strain into JE2 using previously described techniques^63,64^. Bacterial strains used in this study are presented in Supplementary Table S1.

A strain harboring an in-frame deletion of *gisABCD*-*ggt* was constructed using a previously described allelic exchange methodology for *S. aureus*^65^. One kb upstream of SAUSA300_0200 (*gisA*) and one kb downstream of *ggt* were amplified using primers listed in Supplementary Table S2 and cloned into pKOR1-mcs. pKOR1-mcs was confirmed to have correct 1kb homology sequences by Sanger sequencing. The deletion strain was screened for hemolysis on blood agar plates and displayed WT-like hemolysis.

### Sulfur source dependent proliferation analysis

Chemically defined (PN) medium was prepared as previously described^35,66^. PN medium was supplemented with 5 mg mL^−1^ glucose for this work. Prior to inoculation in PN *S. aureus* was cultured in tryptic soy broth (TSB) overnight, washed with PBS, and resuspended in PN medium to an OD_600_ equal to 1. Round-bottom 96-well plates containing PN with 5 mg mL^−1^ glucose supplemented with the indicated sulfur sources were inoculated with *S. aureus* strains at an initial inoculum of OD_600_ of 0.01. Growth analysis was carried out in a Biotek Epoch 2 plate reader set to 37°C with continuous shaking for the indicated time. PN was modified to test sulfur source dependent proliferation of *S. epidermidis* and *S. aureus* by replacing MgSO_4_ with MgCl_2_ and omitting Met, resulting in PN_mod_. *S. aureus* and *S. epidermidis* growth curves were performed as described above in PN_mod_ supplemented with the indicated sulfur sources for 25 h. Sulfur sources were purchased from Millipore Sigma and GSH solutions were freshly prepared prior to each trial to limit oxidation. Alternatively, to ensure CSSC, GSH, and GSSG were maintained in their respective reduced or oxidized forms, stock solutions were prepared by weighing the appropriate amount of the chemical aerobically and immediately transferring it to an anaerobic chamber (Coy) with a 95%:5% nitrogen:hydrogen atmosphere. Sulfur sources were then resuspended in either anaerobically acclimated water (GSH and GSSG) or anaerobically acclimated 1 N HCl (CSSC). Anaerobic proliferation was monitored with the Coy chamber using a Biotek Epoch 2 plate reader.

### Isolation and growth phenotypes of clinical isolates

Four clinical isolates of *S. aureus* were obtained from de-identified specimens at a regional hospital clinical laboratory. Three abscess isolates were confirmed to be methicillin-resistant (strains 1055-1057) and the other was a methicillin-sensitive bone isolate (strain 1059). Identification and minimum inhibitory concentration assays were performed following Clinical and Laboratory Standards Institute approved methods. After initial isolation, subcultures were grown on tryptic soy agar overnight.

### Construction of pET28b∷*ggt* and pET28b∷ *gisA* and purification of tagged protein

*ggt* and *gisA* open reading frames prior to the stop codon were amplified with primers sets listed in Table S2. Expression vector pET28-b was digested with NcoI-HF and XhoI-HF. Plasmid assembly was performed using Gibson assembly and the NEB HiFi assembly kit (NEB, New England, MA). The assembly mixture was transformed into *E. coli*, cells were recovered in lysogeny broth (LB), and plated onto LB agar containing 50 mg mL^−1^ kanamycin and 5 mg mL^−1^ glucose. Plasmids were confirmed using Sanger sequencing and transformed into a NEB strain 3016 *E. coli slyD* mutant^68^. Transformed *E. coli* were cultured in LB with 50 mg mL^−1^ kanamycin overnight at 37°C with shaking at 225 rpm, sub-cultured 1:50 into 500 mL LB with 50 mg mL^−1^ kanamycin in a 2 L flask and grown to an OD_600_ of 0.4-0.7. Ggt or GisA protein expression was induced by addition of 200 μM isopropyl-1-thio-β-D-galactopyranoside (IPTG) and the culture was separated into five 500 mL flasks containing 100 mL of culture and incubated for 4 h at 27°C and 225 rpm shaking. After induction, cells were centrifuged at 10,000 x g for 10 min at 4°C and washed with PBS. Resulting GisA and Ggt induction pellets were resuspended in 40 mL of buffer containing 50 mM tris, 200 mM KCl, 20 mM imidazole at pH 8, or 40-mL buffer containing 50 mM tris, 500 mM NaCl, 20 mM imidazole at pH 8, respectively. Cells were lysed using a fluidizer set to 20,000 psi and samples were run through 5 times. Lysates were then centrifuged at 15,000 x g for 15 min to remove intact cells and the resulting supernatant was retained. To purify the target proteins, Ni-NTA chromatography was used. Purification was performed by incubating the cleared lysate with 1 mL Ni-NTA resin (Qiagen, Hilden, Germany) on a rotating platform at 4°C for 2 h. Protein was eluted with 50 mM tris 400 mM imidazole. Buffers used to purify GisA contained 200 mM KCl while buffers used to purify Ggt contained 500 mM NaCl. Each buffer contained 1x protease inhibitor cocktail (Millipore-Sigma). The GisA elutant was dialyzed using 10 mM tris, 200 mM KCl at pH 7.5 as the dialysis buffer for 18 h. The Ggt elutant was dialyzed using 10 mM tris, 150 mM at pH 7.0 as the dialysis buffer for 18 h. Both the elutions were concentrated using 10 kDa molecular weight cutoff protein concentrators. Purification was confirmed via electrophoresis using 12% SDS-PAGE gels. Protein concentrations were determined with the bicinchoninic acid (BCA) protein kit (BioRAD).

### Quantitation of Ggt enzyme kinetics

Reactions were set up as follows: 10 mM tris with 150 mM NaCl containing 5 μg recombinant Ggt and indicated concentrations of GSH and GSSG dissolved in the reaction buffer. Reactions proceeded for 30 min at 37°C after which samples were incubated at 80°C for 5 min to stop the reaction. Samples were dried using a roto-vac speed vacuum and stored at −80°C until they were hydrated via resuspension in water, derivatized with carboxybenzyl (CBZ), and applied to a Waters Xevo TQ-S triple quadrupole mass spectrometer as previously described^69^. Peak processing was performed by MAVEN, and the signal was normalized to a ^13^C-glutamine internal standard^70^. An external glutamate standard curve was generated using the same chromatographic conditions, and the signal was normalized to a ^13^C-glutamine internal standard. A fit equation to the standard curve was employed to quantitate glutamate within the samples. Glutamate released per min was calculated and data were fit to the Michaelis-Menten equation using GraphPad Prism. Data is the average of glutamate quantified from four independent protein purifications.

### Western blot analysis of *S. aureus* Ggt

The *ggt* ORF, his tag, and stop codon were amplified from pET28b∷*ggt* and cloned into pOS1 P_*lgt*_ digested with NdeI and HindIII using Gibson assembly to generate Ggt-His. Additionally, the *ggt* ORF was amplified from JE2 and pOS1 P_*lgt*_ digested with NdeI and HindIII using Gibson assembly to generate the untagged Ggt. Plasmids were confirmed by Sanger sequencing and transformed from *E. coli* DH5α into *S. aureus* RN4220 via electroporation. Plasmids were purified from RN4220 and transformed into JE2 *ggt*∷Tn. An empty vector control strain was generated by transforming JE2 and *ggt*∷Tn with pOS1 P_*lgt*_. To assess Ggt localization, *S. aureus ggt*∷Tn pOS1 P_*lgt*_∷*ggt* and *ggt*∷Tn pOS1 P_*lgt*_∷*ggt*-His cultures were prepared as described in the proliferation analysis section and sub-cultured into 3, 250 mL flasks each containing 100 mL PN with 25 μM GSSG and 10 μg mL^−1^ chloramphenicol at a starting OD_600_ equal to 0.1. Cells were cultured 4 h at 37°C and 225 rpm shaking. At this time cells reached mid-exponential phase and were recovered via centrifugation, the supernatant retained, and the pellet washed with PBS. Fifty mL of the initial supernatant was precipitated with trichloracetic acid (final percent of 10 % v/v) (TCA), incubated 1 h to overnight at 4°C, pelleted, and the resulting pellet washed twice with 95 % ethanol. Samples were mixed with Laemmli buffer, boiled for 10 min, run on a 12% SDS-PAGE gel using tris-glycine running buffer, and transferred at 65 volts for 1 h at 4°C to PVDF membrane. Forty μg were loaded for WCL, PG, and C+M. Membranes were incubated overnight in phosphate buffered saline tween-20 (PBST) with 3% bovine serum albumin (BSA) at 4°C with agitation. An α-His mouse antibody was used as the primary antibody at a 1:4,000 dilution in PBST supplemented with 5% BSA and incubated for 1 h with shaking. The membrane was washed thrice with PBST. An α-mouse IgG conjugated to horseradish peroxidase (HRP) was used as the secondary antibody at a dilution of 1:4,000 (Sigma-Aldrich). To assess cell wall fractionation a mouse α-protein A (Spa) primary antibody was used at 1:6,000 dilution. Membranes were washed thrice in PBST after incubation with secondary antibody. Membranes were developed using the ECL Prime kit (Cytiva, Marlborough, MA) and imaged using Amersham Imager 600 (GE Healthcare, Amersham, Buckinghamshire, UK). Fractionation was repeated with 3 independent sets of cultures and one set of immunoblots is presented as a representative.

### Identification of GisABCD-Ggt homologues across bacteria

The USA300_FPR3757 (assembly GCF_000013465.1) Ggt protein sequence (ABD22038.1) was used as the query protein for homology searches (using DELTA-BLAST) using the NCBI RefSeq database^69–71^. Data was filtered to include only Firmicutes that encoded Ggt homologues containing a glutamyl transpeptidase domain. This dataset was used as the subject to query USA300_FPR3757 GisABCD using DELTA-BLAST. Percent similarities to the *S. aureus* GisABCD-Ggt protein amino acid sequences were used to generate a heatmap. The heatmap and hierarchical clustering of similar protein profiles were generated using the R package, pheatmap.

### Quantitation of *S. aureus* and *S. epidermidis* competition

*S. aureus, S. aureus ΔgisABCD-ggt*, and *S. epidermidis* were cultured overnight in TSB, pelleted, wash in PBS, and normalized to the same OD_600_ in PN_mod_. Strains were mixed in a 1:1 ratio (v/v) and inoculated into 5 mL PN_mod_. PN_mod_ was supplemented with 25 μM GSSG, 50 μM GSH, 750 μM GSH, or 50 μM Met. Dilution plating of the freshly mixed co-culture was plated onto mannitol salt agar (MSA) to quantify initial counts of each organism. Cultures were incubated for 24 h at 37°C with 225 rpm shaking after which the cultures were dilution plated onto MSA and allowed to grow for 48 h at 35°C. *S. aureus* ferments mannitol and appears yellow on MSA, while *S. epidermidis* does not and maintains a pink color; consequently, yellow and pink colored colonies were enumerated to assess quantities of each organism. Competitive indices (CI) were calculated by dividing the *S. aureus* to *S. epidermidis* output ratio by the *S. aureus* to *S. epidermidis* input ratio. A CI greater than one indicates more *S. aureus* than *S. epidermidis* while a CI less than one signifies greater quantities of *S. epidermidis* compared to *S. aureus*.

## Acknowledgements

The defined transposon mutant library used in this study was provided by the Network on Antimicrobial Resistance in *Staphylococcus aureus* (NARSA) for distribution by BEI Resources, NIAID, NIH: Nebraska Transposon Mutant Library (NTML) Screening Array NR-48501. We thank the laboratory of Dr. Taeok Bae at Indiana University for supplying the pKOR1-mcs plasmid, Dr. Anthony Richardson for the *S. epidermidis* strain RP62a, and we thank the Dr. Victor DiRita and Dr. Sean Crosson laboratories at Michigan State University for technical support. We thank Dr. Martin P. Ogrodzinski and the MSU Mass Spectrometry and Metabolomics Core for technical support. This work is funded by the National Institutes of Health R01 AI139074 and R21 AI142517.

## Notes

### Competing Interest Statement

The authors have declared no competing interest.

