## Supplemental Information for "The glutathione import system satisfies the *Staphylococcus aureus* nutrient sulfur requirement and promotes interspecies competition"

**Supplementary Information**

**Supplementary Materials and Methods**

**Domain prediction of GisABCD-Ggt.** The USA300_FPR3757 (assembly GCF_000013465.1) reference was used to predict domain architectures for GisA (ABD21741.1), GisB (ABD21022.1) GisC (ABD20640.1), GisD (ABD22752.1), and Ggt (ABD22038.1). Protein sequences were analyzed with custom scripts using InterProScan, TMHMM, Phobius, Pfam, and PROSITE to identify domains, secondary structures, and cellular localization signatures[1–6]. Domain architectures were visualized using custom R scripts and the R package, gggenes[7].

**GisA ATPase activity assay.** ATPase activity of purified recombinant GisA was monitored using the Malachite Green Phosphate assay (Millipore-Sigma). Recombinant GisA was diluted to 1 µg per reaction. Diluted GisA, 250 µM $MgCl_2$, and 400 µM ATP were incubated for 1 h at 37°C and samples were taken at 0, 15, 30, 45, and 60 min. At the indicated time points samples were flash frozen in a dry-ice ethanol bath and stored at -80°C[8]. Samples were thawed at room temperature, Malachite Green reagent was added, and $P_i$ release was determined following the manufacturer's instructions. Reactions containing ATP in the absence of GisA were used to correct absorbance measurements due to residual inorganic phosphate and non-enzymatic ATP hydrolysis. Each biological replicate used an independently purified GisA preparation.

**Murine systemic infections.** WT and *gisB*::Tn were cultured in TSB overnight at 37°C, diluted 1:100 into TSB, and cultured for 3 h at 37°C at 225 rpm shaking. Cultures were pelleted, washed with PBS and normalized to $OD_{600}$ equal to 0.4. Female C57BL6 mice were retro-orbitally infected with $10^7$ CFUs and infection proceeded for 96 h after which heart, liver and kidneys were collected and homogenized in 1 mL PBS. Organ homogenates were serially diluted and plated onto TSA. Bacterial burdens quantified as CFUs $mL^{-1}$ were determined. Infections were performed at

Michigan State University under the principles and guidelines described in the Guide for the Care and Use of Laboratory Animals[9]. Animal work was followed as approved by Michigan State University Institutional Animal Care and Use Committee (IACUC) approved protocol number 12/16-205-00.

**Supplementary Table S1. *Staphylococcus* strains used in this study.**

| strain | description | reference |
|---|---|---|
| **methicillin-resistant *S. aureus*** | | |
| JE2 | Laboratory derived wild type parental MRSA; USA300_LAC; CC8 | 10 |
| MW2 | MRSA, USA400; CC1 | 11,12 |
| COL | MRSA; CC8 | 13,14 |
| SF8300 | CA-MRSA; USA300; CC8 | 15 |
| TCH1516 | MRSA; USA300; CC8 | ATCC |
| SAUSA300_0200::Tn | NTML NE392 Tn insertion in *gisA* | 10 |
| SAUSA300_0201::Tn | NTML NE541 Tn insertion in *gisB* | 10 |
| SAUSA300_0202::Tn | NTML NE457 Tn insertion in *gisC* | 10 |
| SAUSA300_0203::Tn | NTML NE215 Tn insertion in *gisD* | 10 |
| *ggt*::Tn | NTML NE254 Tn insertion in *ggt* | 10 |
| *gisA*::Tn | *gisA* mutant strain backcrossed into JE2 | this study |
| *gisB*::Tn | *gisB* mutant strain backcrossed into JE2 | this study |
| *gisC*::Tn | *gisC* mutant strain backcrossed into JE2 | this study |
| *gisD*::Tn | *gisD* mutant strain backcrossed into JE2 | this study |
| *ggt*::Tn | *ggt* mutant strain backcrossed into JE2 | this study |
| Δ*gisABCD-ggt* | in-frame deletion of *gisABCD-ggt* in JE2 | this study |
| JE2 pOS1 $P_{lgt}$ | JE2 harboring pOS1 $P_{lgt}$ empty vector | this study |
| *ggt*::Tn pOS1 $P_{lgt}$ | backcrossed JE2 *ggt*::Tn harboring pOS1 $P_{lgt}$ empty vector | this study |
| *ggt::*Tn pOS1 $P_{lgt}$::*ggt* | backcrossed JE2 *ggt*::Tn harboring pOS1 $P_{lgt}$::*ggt* | this study |
| *ggt*::Tn pOS1 $P_{lgt}$::*ggt*-His | backcrossed JE2 *ggt*::Tn harboring pOS1 $P_{lgt}$::*ggt* encoding a His-tag | this study |
| **clinical isolates** | | |
| 1055 | MRSA abscess hand cellulitis | this study |
| 1056 | MRSA abscess left arm | this study |
| 1057 | MRSA left wrist/ index finger | this study |
| 1059 | MSSA bone from the coccyx/chronic osteomyelitis | this study |
| ***Staphylococcus epidermidis* strains** | | |
| *Staphylococcus epidermidis* | strain RP62a | 16 |
| *Staphylococcus epidermidis* | clinical isolate | this study |

**Supplementary Table S2. Primers used in this study.**

| name | sequence 5’-3’ | description |
|---|---|---|
| pET28b::*ggt* F | AAGAAGGAGATATACCATGGTCATTAACTTA AATGACAAAC | amplify *ggt* ORF without stop codon to clone into pET28b |
| pET28b::*ggt* R | GATGATGGCTGCTGCTGCCCATGTCTTGTG ATACTATCTCGAT | amplify *ggt* ORF without stop codon to clone into pET28b |
| pKOR1-mcs Δ*gis* upstream F | CTGCTAGCTAGCTAGAGATATCAAACGATAA AAAATATACAAATAAAAATCTAATTGTAG | amplify 1kB upstream of SAUSA300_0201 to clone into pKOR1-mcs |
| pKOR1 Δ*gis* upstream R | AGCGTATAAAAAGTCATGCGTTGTGCAAC | amplify 1 kB upstream of SAUSA300_0201 to clone into pKOR1-mcs |
| pKOR1-mcs Δgis downstream F | CGCATGACTTTTTATACGCTTGATATGAAGT TTG | amplify 1 kB downstream of SASUA300_0204 to clone into pKOR1-mcs |
| pKOR1-mcs Δgis downstream R | CGG AAC CGG TAC CAA TGG ATA TCT ATG TTT TTG GCA ATG AAG TG | amplify 1kB downstream of SAUSA300_0204 to clone into pKOR1-mcs |
| Δ*gis* conf. F | GACTAAGCTAAGTTGACACAC | confirmation of gis deletion |
| Δ*gis* conf. R | CATCCAAATCATCTATTAAAATCC | confirmation of gis deletion |
| pET28b::*gisA* F | ACTTTAAGAAGGAGATATACATGTCAAATTTA TTAGAAGTCAAC | amplify *gisA* ORF without stop codon to clone into pET28b |
| pET28b::*gisA* R | AGTGGTGGTGGTGGTGGTGCGATTTAGCAA TAACTGCTAC | amplify *gisA* ORF without stop codon to clone into pET28b |
| pOS1 $P_{lgt}$::*ggt* F | ACAATTGAGGTGAACATATGGTCATTAACTT AAATGACAAAC | amplify *ggt* ORF to clone into pOS1 $P_{lgt}$ |
| pOS1 $P_{lgt}$::*ggt* R | CTACCCCCTTGTTTGGATCCCTATCTTGTGA TACTATCTC | amplify *ggt* ORF to clone into pOS1 $P_{lgt}$ reverse primer |
| pOS1 $P_{lgt}$::*ggt*-his F | AAATACAATTGAGGTGAACATATGGTCATTA ACTTAAATGACAAACAG | amplify *ggt* with his tag from pET28B::*ggt* to clone into pOS1 $P_{lgt}$ |
| pOS1 $P_{lgt}$::*ggt*-his R | AGCTTGGCTGCAGGTCGACGGATCC TCAGTGGTGGTGGTGGTG | amplify *ggt* with his tag from pET28B::*ggt* to clone into pOS1 $P_{lgt}$ |

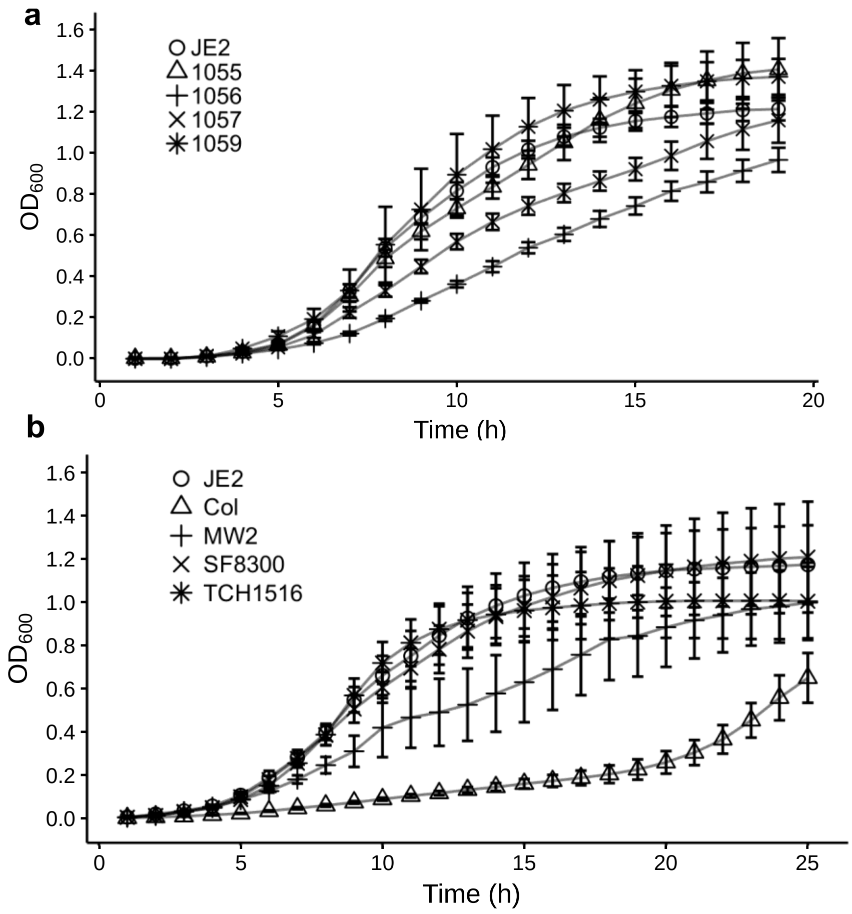


**Supplementary Figure S1. Supplementation with GSSG as the sole source of nutrient sulfur stimulates proliferation of *S. aureus*.** Laboratory derived JE2 and clinical isolates (**a**) or other laboratory strains (**b**) were cultured in medium containing 25 µM GSSG. The mean $OD_{600}$ of three independent trials is depicted and error bars represent ± 1 standard error of the mean.

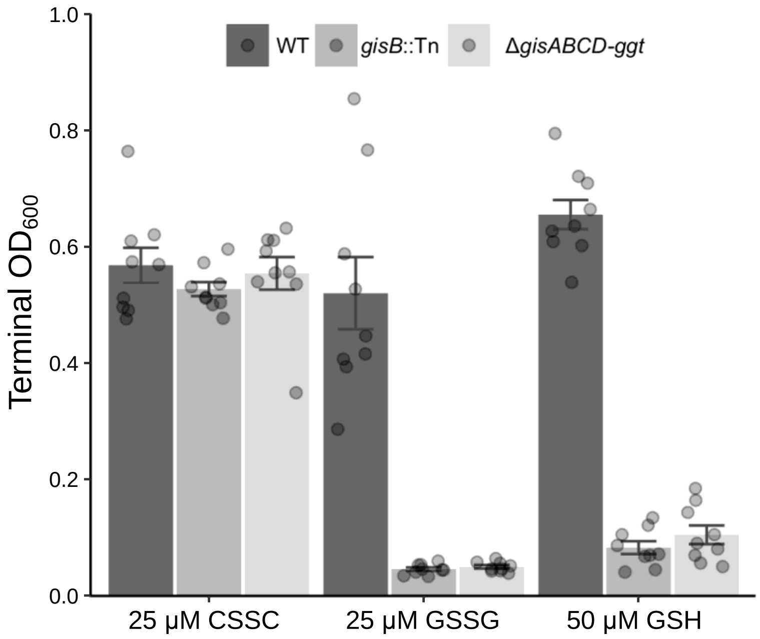


**Supplementary Figure S2. GisABCD-Ggt promotes anaerobic proliferation in PN medium supplemented with GSSG or GSH.** WT, *gisB*::Tn, and Δ*gisABCD-ggt* were cultured in chemically defined PN medium in the presence of the alternative terminal electron acceptor sodium nitrate (100 mM) and the indicated sulfur sources. Sulfur source stock solutions were prepared anaerobically to maintain respective reduced or oxidized states. Each point represents the mean terminal $OD_{600}$ after 24 h of growth derived from a technical triplicate (1 trial) and each bar represents the mean of nine independent trials. Error bars represent ± 1 standard error of the mean.

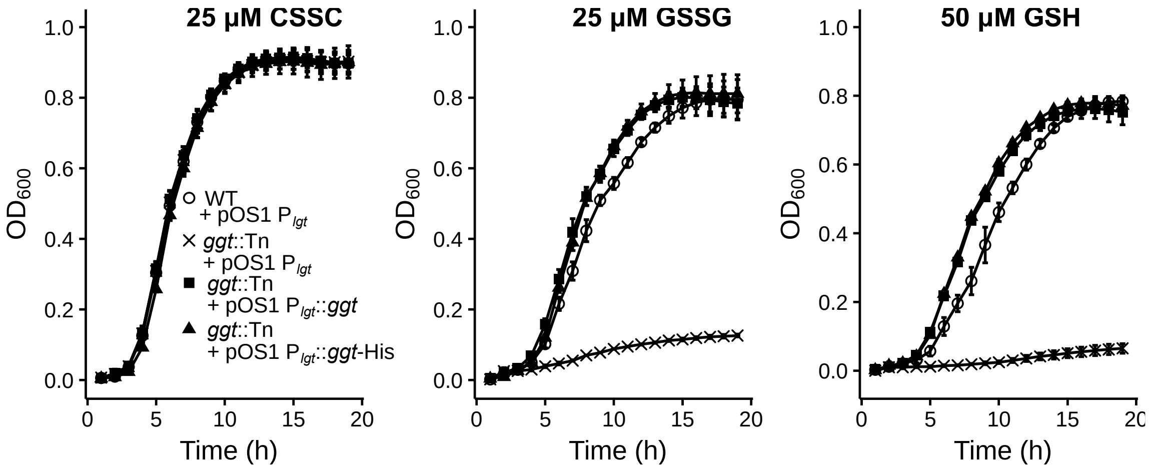


**Supplementary Figure S3. Ectopic expression of native or His-tagged Ggt complements *ggt* mutant proliferation in medium supplemented with reduced or oxidized GSH.** Strains cultured in medium supplemented with 25 µM CSSC, 25 µM GSSG, or 50 µM GSH. Presented is the mean of at least three independent trials and error bars represent ± 1 standard error of the mean.

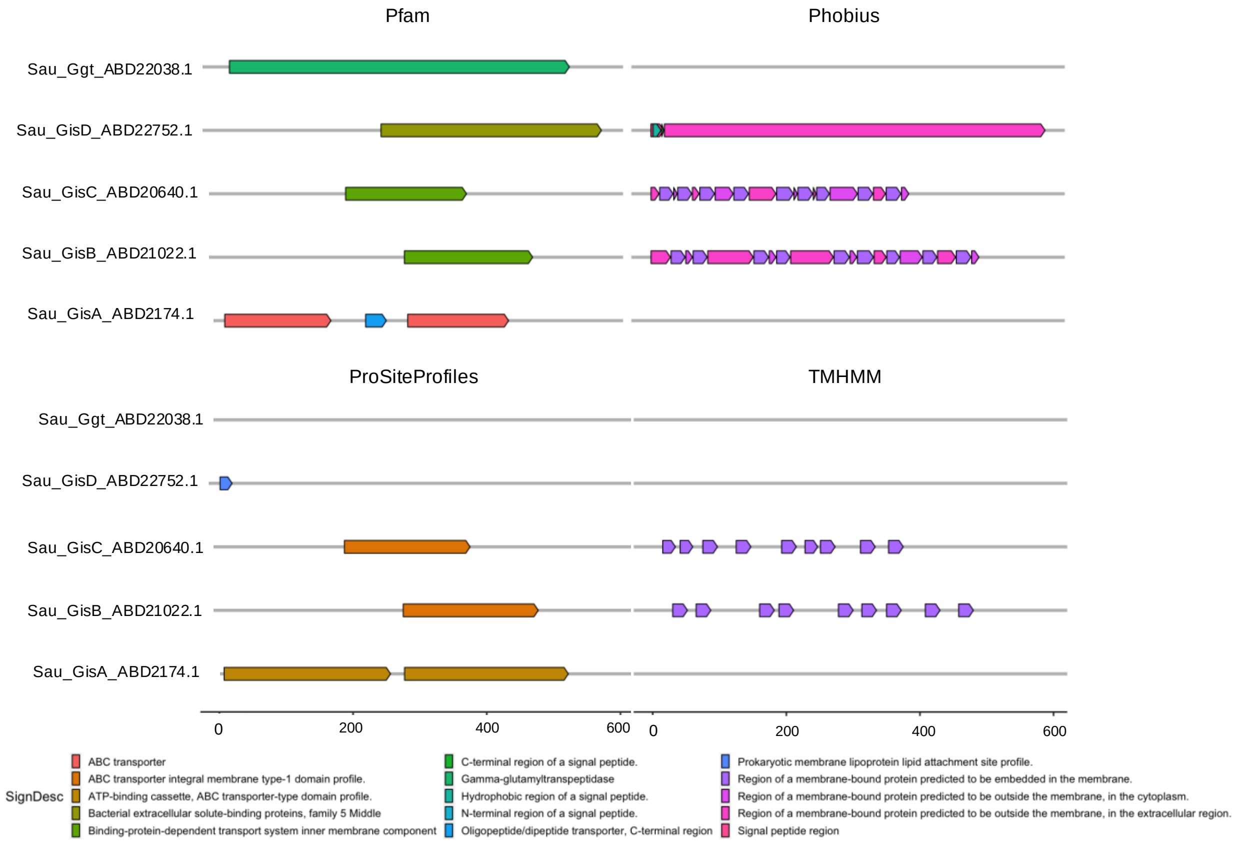


**Supplementary Figure S4. Domain architectures and secondary structure predictions for the *S. aureus* GisABCD-Ggt system.** Domains were predicted using InterProScan[4,5], specifically using profile databases Pfam, ProSiteProfiles, and prediction algorithms, Phobius and TMHMM for the query proteins: GisA (ABD21741.1), GisB (ABD21022.1) GisC (ABD20640.1), GisD (ABD22752.1), and Ggt (ABD22038.1).

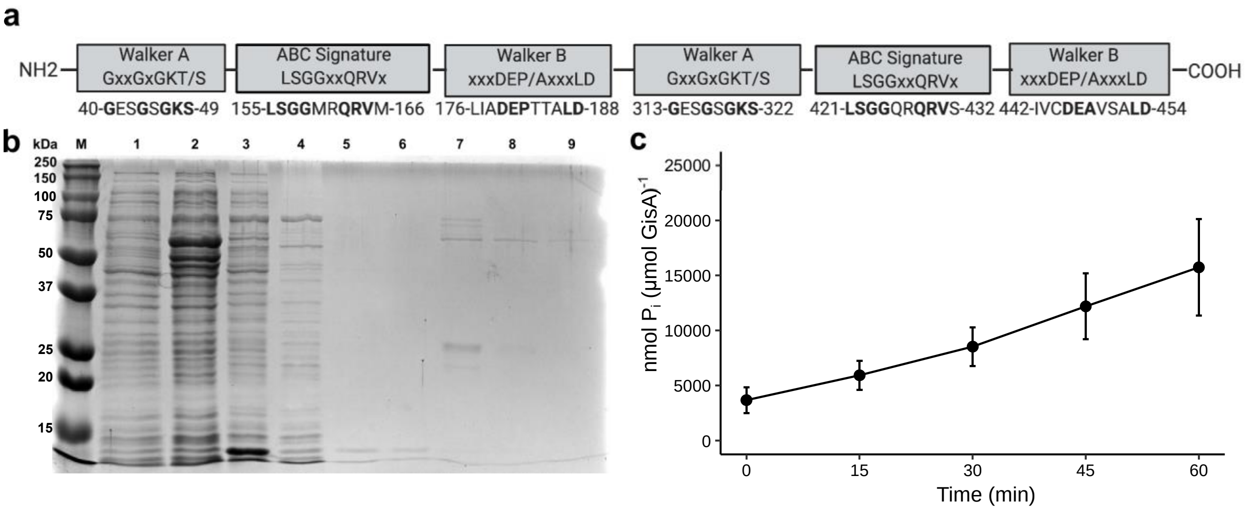


**Supplementary Figure S5. GisA encodes ATPase domain signatures and demonstrates ATP hydrolysis activity. a,** GisA contains two Walker A motifs, two Walker B motifs, and two ABC Signature motifs. Numbers correspond to the codon position of the first amino acid in the motif. Domain structure illustration was created with BioRender. **b,** Recombinant histidine (His)-tagged GisA was expressed and purified from a modified *E. coli* NEB 3016 expression strain. Indicated samples and fractions were collected during expression and subsequent purification. Samples were loaded onto 12% SDS-PAGE and strained with Coomassie blue. Lanes: molecular weight ladder (M); 1: uninduced whole cell lysate (WCL); 2: WCL 4 h post IPTG induction; 3: pre-column lysate; 4: Ni-NTA flow through; 5: and 6: 20 mM imidazole elution; 7: 100 mM imidazole elution; 8: and 9: 400 mM imidazole elution. GisA is predicted to be 59 kDa. **c,** Time course analysis of ATP hydrolysis activity of GisA incubated at 37°C for 1 h with 400 μM ATP. Samples were taken at the indicated time points and inorganic phosphate ($P_i$) concentrations were determined using the malachite green assay. Presented is the mean of nine independent trials and error bars represent ± 1 standard error of the mean. Each trial used a new purification of GisA.

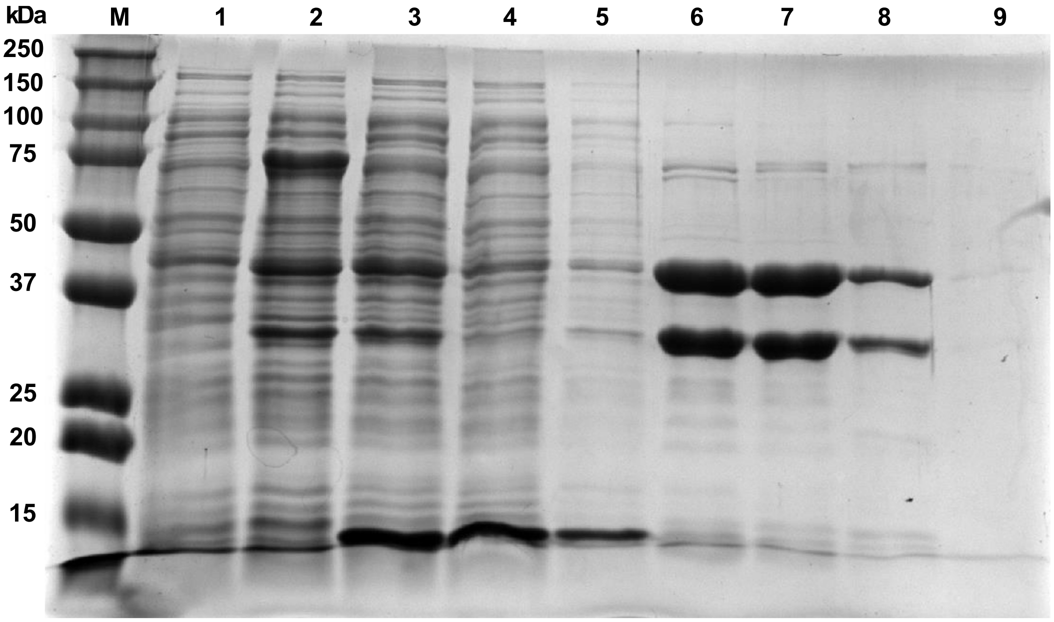


**Supplementary Figure S6. Heterologous expression and purification of *S. aureus* Ggt from *E. coli*.** Recombinant His-tagged Ggt was expressed and purified from a modified *E. coli* NEB 3016 expression strain. Indicated samples and fractions were collected during expression and subsequent purification. Samples were loaded onto 12% SDS-PAGE and strained with Coomassie blue. Lanes: molecular weight ladder (M); 1: uninduced whole cell lysate (WCL); 2; WCL 4 h post IPTG induction; 3, pre-column lysate; 4, Ni-NTA column flow through; 5, 20 mM imidazole wash; 6: 50 mM imidazole elution; 7: 100 mM imidazole elution; fractions 8 and 9: consecutive 400 mM imidazole elutions.

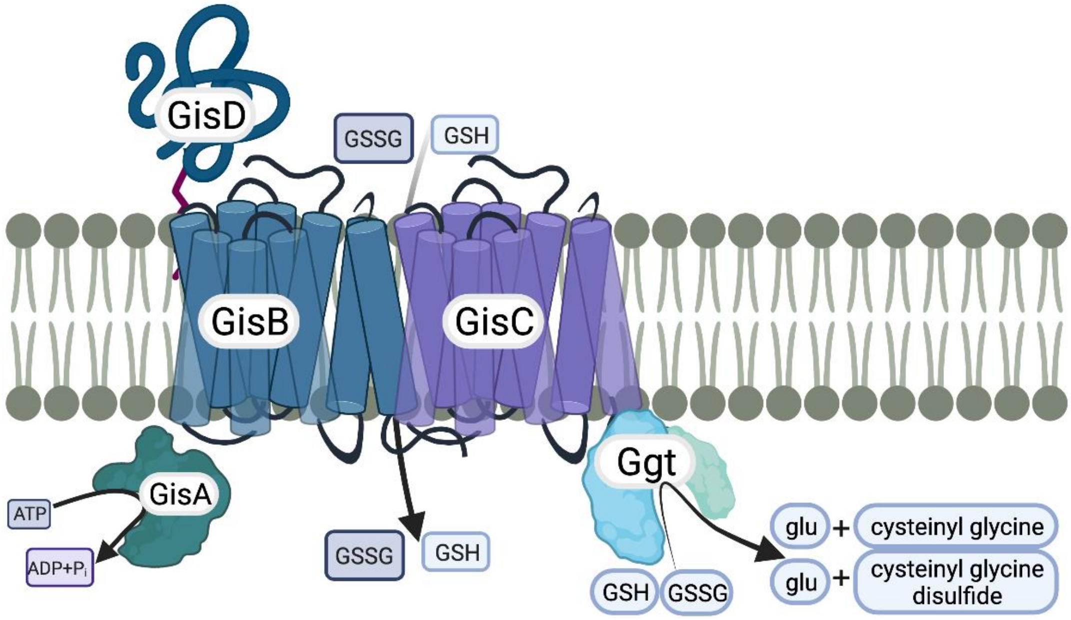


**Supplementary Figure S7. A model of GisABCD-Ggt mediated acquisition of GSH and GSSG.** Bioinformatic predictions and experimental evidence supports the following illustration of *S. aureus* import and catabolism of exogenous GSH and GSSG. GisD, a predicted substrate binding protein, binds GSH or GSSG in the extracellular milieu, which are transported into the cytoplasm by the transmembrane permease complex, GisBC. GisA hydrolysis of ATP provides energy needed for import. Finally, GSH and GSSG are cleaved in the cytoplasm by Ggt, generating glutamate and cysteinyl-glycine or cysteinyl-glycine disulfide, depending on the substrate. The model illustration was created using BioRender. Adapted from membrane proteins by BioRender.com (2021). Retrieved from http://app.biorender.com/biorender-templates.

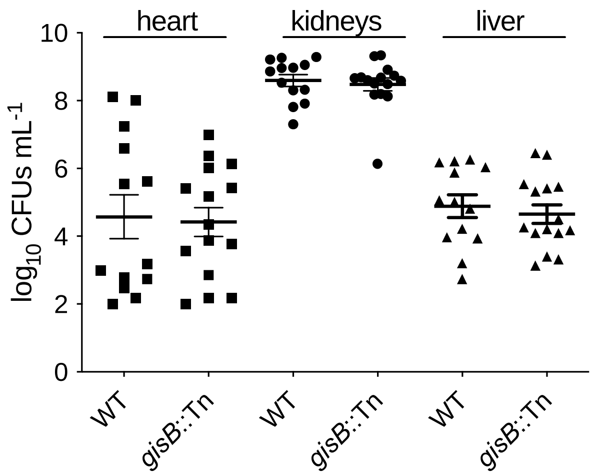


**Supplementary Figure S8. Virulence of the *gisB*::Tn mutant strain mimics wild type.** Bacterial burdens within indicated organs of C57BL/6 mice were enumerated after 96 h of systemic infection with either WT or *gisB*::Tn. Bacterial burdens are presented as $\log_{10}$ CFUs $mL^{-1}$ for liver, combined kidneys, and heart. The line represents the mean and error bars represent ± 1 standard error of the mean.

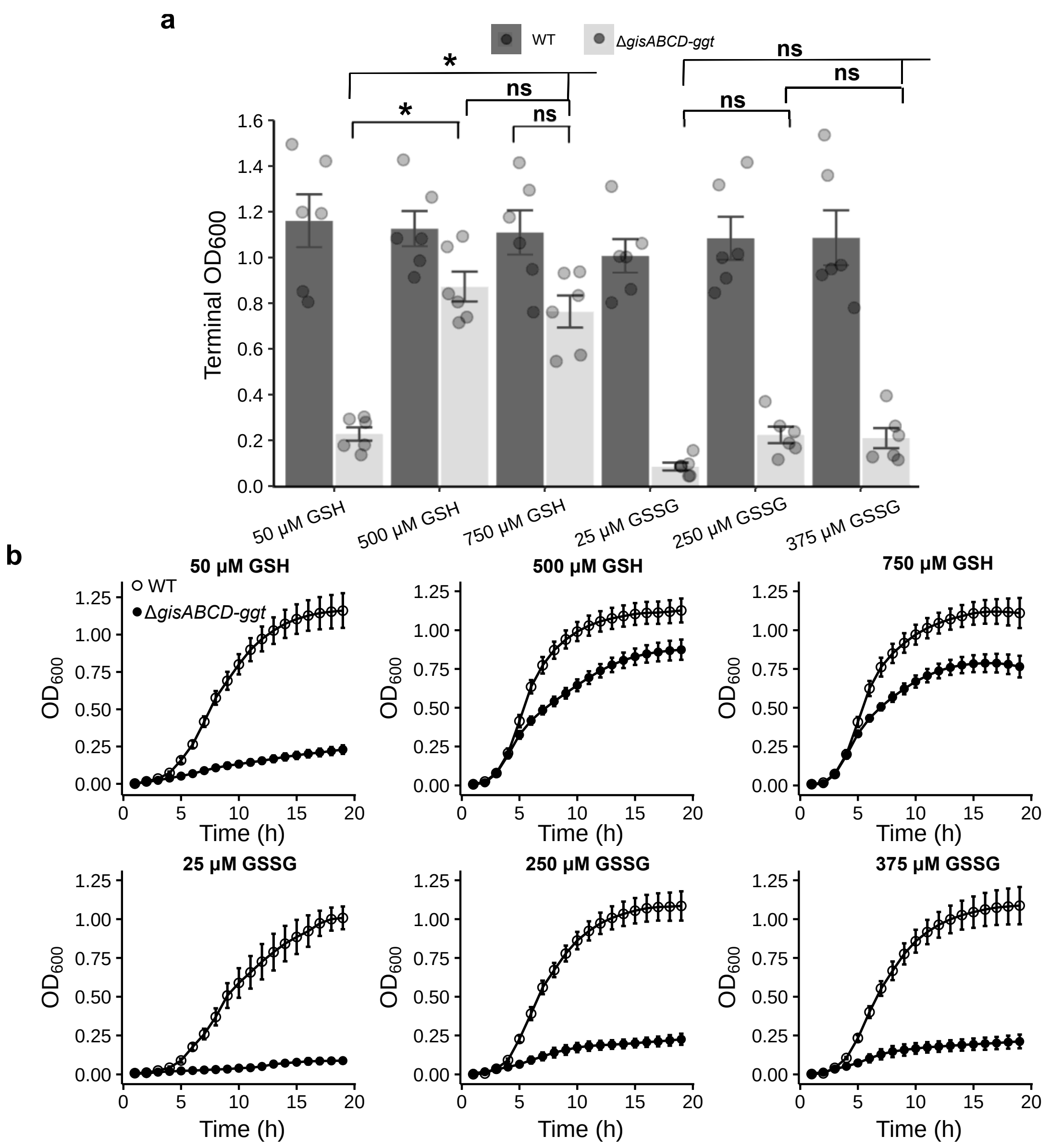


**Supplementary Figure S9.** ***S. aureus*** **acquires GSH independent of GisABCD-Ggt in physiologically relevant concentrations of GSH**. **a,** WT and Δ*gisABCD-ggt* were cultured in medium supplemented with 50 μM GSH, 500 μM GSH, 750 μM GSH, 25 μM GSSG, 250 μM GSSG, or 375 μM GSSG. Bars depict the mean $OD_{600}$ after 19 h of growth. Data points represent the terminal $OD_{600}$ from each individual trial. Error bars represent ± 1 standard error of the mean. ***** Indicates p-value <0.05 by one-way ANOVA with Tukey's multiple comparison correction. **b**, Corresponding growth curves for WT and Δ*gisABCD-ggt* cultured in increasing concentrations of GSH or GSSG.

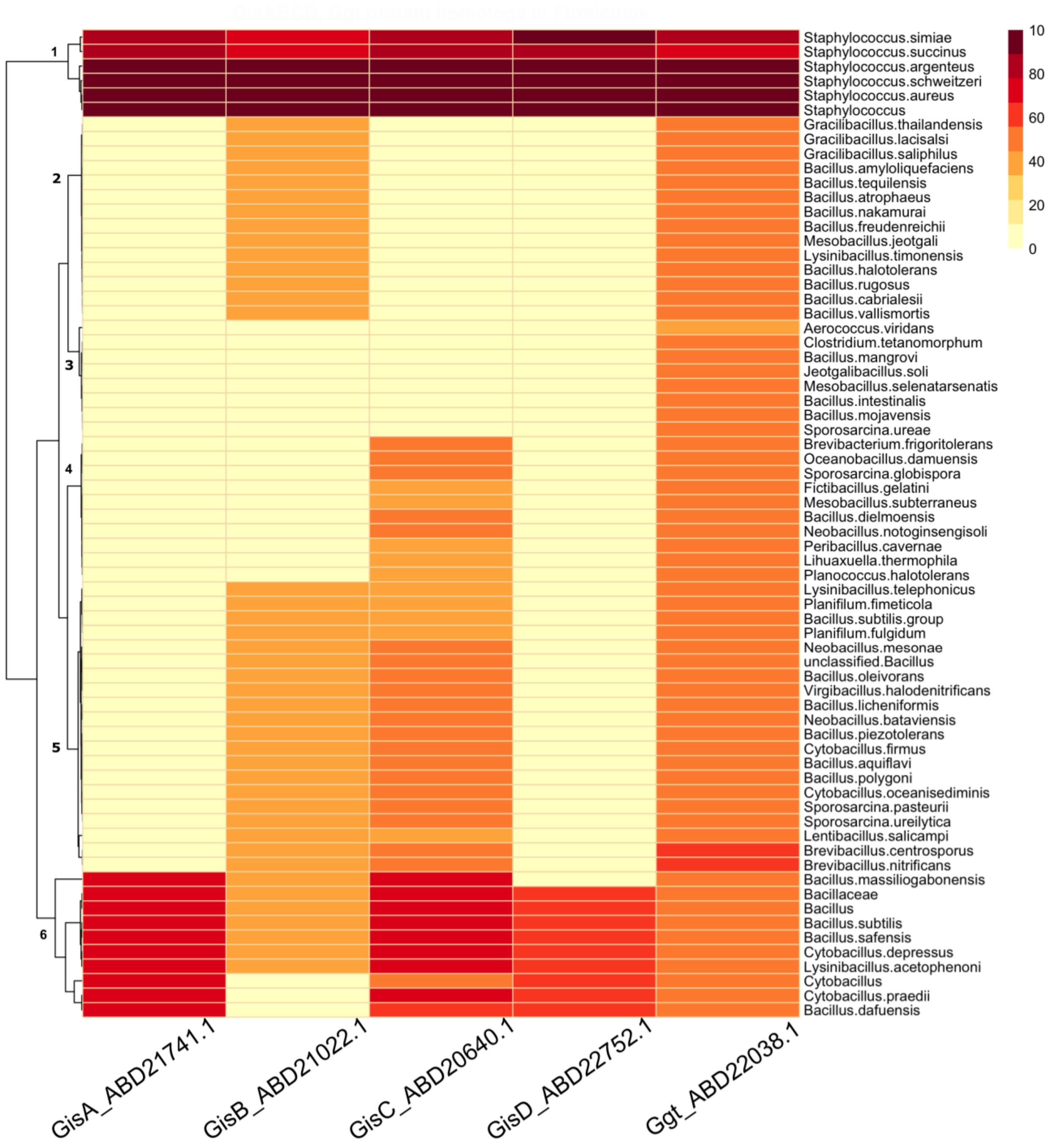


**Supplementary Figure S10. Conservation of Ggt and GisABCD across Firmicutes.** Percent similarity of *S. aureus* Ggt was queried, and results were limited to Firmicutes encoding proteins that harbor an annotated γ-glutamyl transpeptidase domain. *S. aureus* GisABCD was subsequently queried using this dataset. The color of the box corresponds to the percent similarity to *S. aureus* Ggt. Hierarchical clustering is presented in the dendrogram on the left.

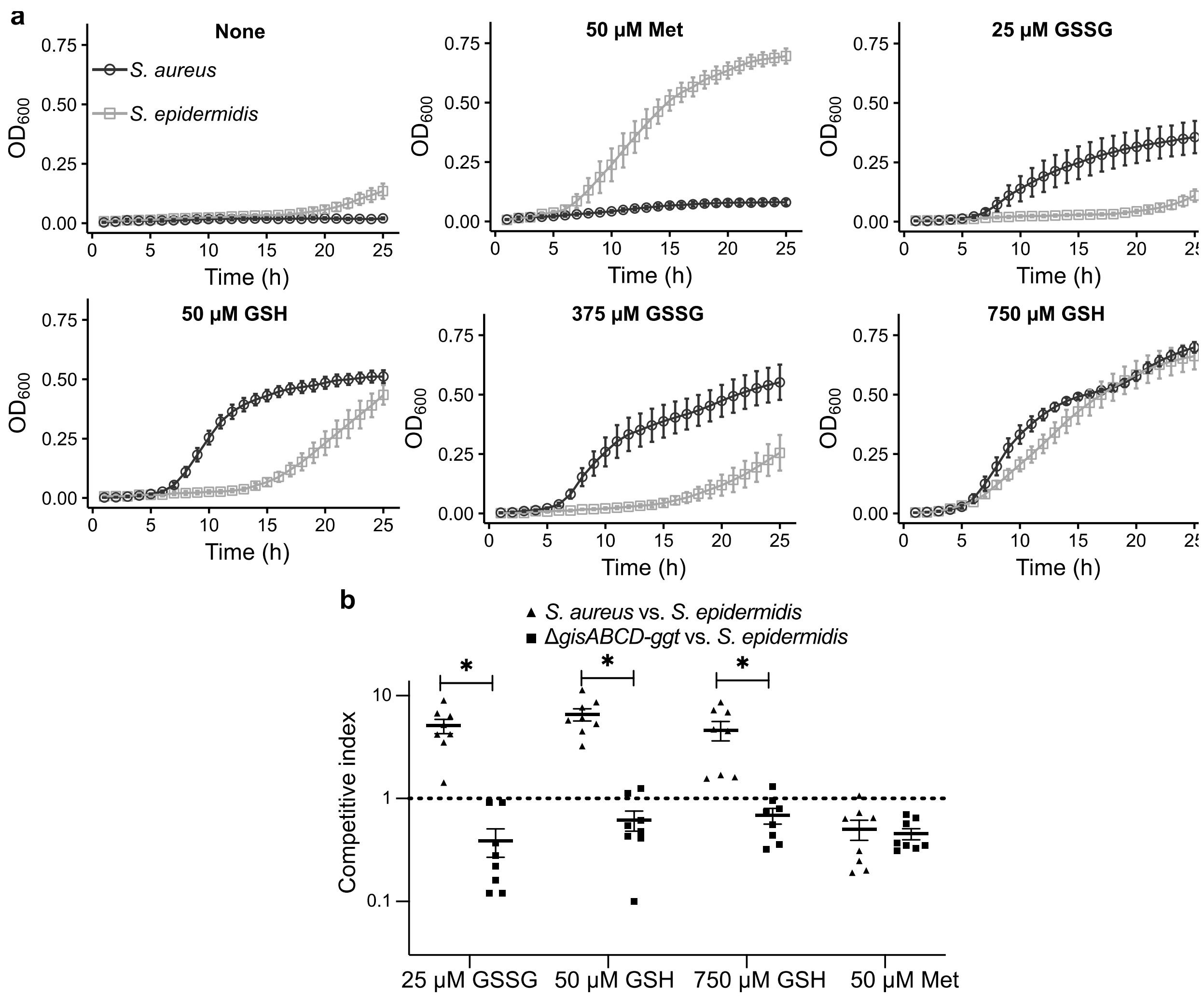


**Supplementary Figure S11. *S. epidermidis* and *S. aureus* nutrient sulfur source utilization is distinct and promotes interspecies competition. a,** *S. aureus* and *S. epidermidis* were cultured in $PN_{mod}$ containing the indicated source of nutrient sulfur. The mean $OD_{600}$ of at least three independent trials and the error bars depict ± 1 standard error of the mean are presented. **b**, *In vitro* competition between *S. epidermidis* and *S. aureus* in $PN_{mod}$ containing the indicted sources of nutrient sulfur. The competitive index of each trial is presented. The line represents the mean and the error bars depict ± 1 standard error of the mean. ***** Indicates p-value <0.05 by one-way ANOVA with Tukey's multiple comparison correction.
